# CompHEAR: A Customizable and Scalable Web-Enabled Auditory Performance Evaluation Platform for Cochlear Implant Sound Processing Research

**DOI:** 10.1101/2023.12.22.573126

**Authors:** Kris Merrill, Leah Muller, Jordan A. Beim, Phillipp Hehrmann, Dean Swan, Daniel Alfsmann, Tony Spahr, Leo Litvak, Andrew J. Oxenham, Aaron D. Tward

**Author notes:** All correspondence should be addressed to: Aaron Tward, Department of Otolaryngology-Head and Neck Surgery, University of California, San Francisco, 513 Parnassus Avenue, HSE 740, San Francisco, CA, USA.

## Abstract

**Objective:** Cochlear implants (CIs) are auditory prostheses for individuals with severe to profound hearing loss, offering substantial but incomplete restoration of hearing function by stimulating the auditory nerve using electrodes. However, progress in CI performance and innovation has been constrained by the inability to rapidly test multiple sound processing strategies. Current research interfaces provided by major CI manufacturers have limitations in supporting a wide range of auditory experiments due to portability, programming difficulties, and the lack of direct comparison between sound processing algorithms. To address these limitations, we present the CompHEAR research platform, designed specifically for the Cochlear Implant Hackathon, enabling researchers to conduct diverse auditory experiments on a large scale.

**Study Design:** Quasi-experimental

**Setting:** Virtual

**Methods:** CompHEAR is an open-source, user-friendly platform which offers flexibility and ease of customization, allowing researchers to set up a broad set of auditory experiments. CompHEAR employs a vocoder to simulate novel sound coding strategies for CIs. It facilitates even distribution of listening tasks among participants and delivers real-time metrics for evaluation. The software architecture underlies the platform’s flexibility in experimental design and its wide range of applications in sound processing research.

**Results:** Performance testing of the CompHEAR platform ensured that it could support at least 10,000 concurrent users. The CompHEAR platform was successfully implemented during the COVID-19 pandemic and enabled global collaboration for the CI Hackathon (www.cihackathon.com).

**Conclusion:** The CompHEAR platform is a useful research tool that permits comparing diverse signal processing strategies across a variety of auditory tasks with crowdsourced judging. Its versatility, scalability, and ease of use can enable further research with the goal of promoting advancements in cochlear implant performance and improved patient outcomes.

## Introduction

Cochlear implants (CIs) are auditory prostheses that can partially restore hearing to those with severe to profound hearing loss by directly stimulating the auditory nerve through electrodes. However, despite four decades of availability, recent years have seen modest improvements in CI performance. In particular, CI recipients often struggle with sound clarity, understanding speech in noisy environments, and music appreciation.(1,2) This raises an important question: have we reached the peak performance of CI hardware, or is there still untapped potential for significant improvements in sound processing strategies through innovative approaches?

Research hardware and software platforms and interfaces have been crucial for advancements in hearing device research(3–5). However, existing research interfaces provided by major CI manufacturers are limited in supporting a wide range of auditory experiments due to issues related to portability and ease of programming. Moreover, reliable evaluation of auditory performance in CI recipients typically require a sound-proof room with specialized and well calibrated equipment to ensure accurate measurements and to eliminate external auditory distractions. Audiologists interpret hearing performance using a standardized battery of auditory stimuli designed to evaluate abilities in various listening conditions which typically consists of prerecorded monosyllabic words, sentences in quiet, and speech in noise components with challenging signal-to-noise ratios. Because of challenges related to clinical testing of novel technologies in CI users, an alternative method to evaluate sound processing strategies relies on vocoder simulations of processed sound signals to predict outcomes in CI users.(6–10) Although vocoded simulations are challenging to broadly validate against the true experience of hearing with a CI, they provide a useful way to compare many different parameters that would be challenging in real CI users. However, because of the diversity of approaches used by various groups, independently generated strategies are not always directly compared in a standardized manner.

To overcome these constraints and foster innovation in CI sound processing, the Cochlear Implant Hackathon emerged as a collaborative effort between Advanced Bionics, the University of California San Francisco, and the University of Minnesota. The Hackathon, conducted entirely online from December 2020 to January 2021, aimed to inspire the public to improve CI sound processing strategies. The virtual competition format encouraged international collaboration and enabled entrants and judges to participate from anywhere with internet access.

The primary objective of the Cochlear Implant Hackathon was to generate novel ideas for signal-processing strategies to enhance CI sound quality for users. Our secondary goal was to prototype a software platform, accessible to researchers and the public, for standardizing direct comparison of different CI sound processing strategies across various auditory tasks for use in future efforts to improve sound processing strategies. To facilitate this, we developed a user-friendly and scalable platform, which enabled customized listening experiments with minimal training. The platform ensured equal distribution of tasks across participants and real-time interpretation of metrics thus enabling researchers to conduct comprehensive evaluations of sound processing algorithms remotely and at scale.

This study presents a high-level overview of the CompHEAR platform, which was designed to enable the CI Hackathon (cihackathon.com) and implemented to foster innovation in CI sound processing. Through scalable and customizable evaluation methods, researchers can confidently rank sound processing strategies across various auditory categories. This approach offers reliable and thorough evaluations, with the ultimate goal of leading to improved auditory performance for CI users.

### Cochlear Implant Hackathon

The Cochlear Implant Hackathon was a dynamic competition facilitated by our custom-built research platform, illustrated in Figure 1. The platform supported both participants (hackers) who submitted innovative sound processing algorithms and judges who volunteered to evaluate them. Hackers were equipped with resources, including Advanced Bionics’ fully featured CI sound coding strategy, a vocoder CI simulation framework, and sample audio files. Through the platform, participants submitted entries, which were then transformed into CI simulation audio samples. These simulations were randomly distributed among multiple sets of auditory tasks for multiple rounds of evaluation. Organizers had real-time access to scores as entries progressed through the crowdsourced judging process, and winners received prizes. The CI hackathon website can be visited at https://cihackathon.com.

**Figure 1.**
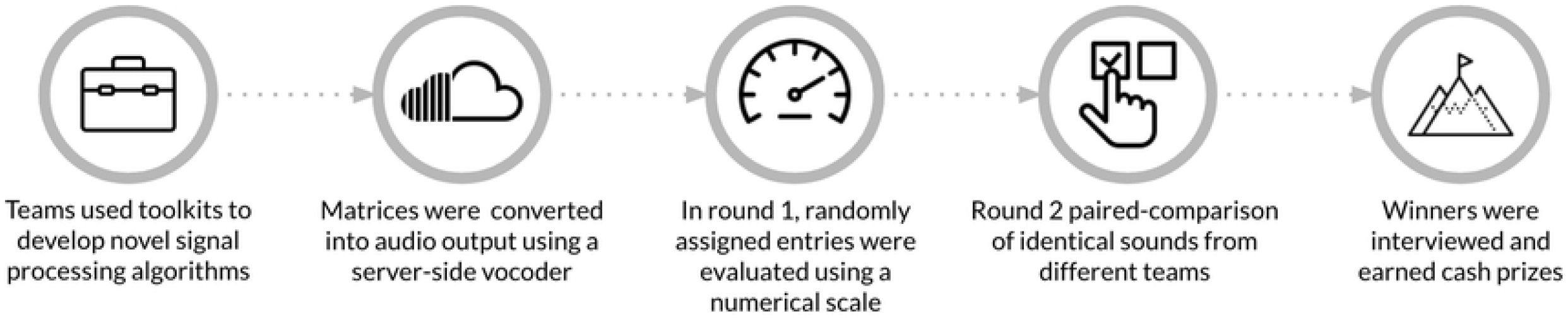
Schematic overview of the Cochlear Implant Hackathon.

### CompHEAR Research Platform

The source code for the CompHEAR platform is available on GitHub at: https://github.com/kmerri11/cihackathon-public. Our web-based sound processing evaluation platform demonstrated scalability and flexibility, allowing customization to meet the needs of researchers. It was developed and hosted on Amazon Web Services (AWS), utilizing various AWS technologies. The platform featured secure logins and validation for uploaded submissions, a backend application interface for a vocoder, and customizable evaluation frameworks for judging. An administrative interface provided easy monitoring of team progress, individual judges’ progress, and the status of each evaluation. A real-time leaderboard displayed the results. The platform was developed using open-source libraries in NodeJS and Python. Entries and processed auditory CI simulations were stored in AWS S3, an enterprise cloud storage system. DynamoDB, a key-value storage system, was used to track entries and collect data. AWS Lambda served web requests, allowing virtually unlimited scaling. GitHub was utilized as our version control system and source code repository, with automated software deployments triggered by code changes. Before public release, a separate staging site was used for quality checks.

#### Input sound files

The research platform was specifically designed to support auditory input in the form of wave (.wav) files. For the hackathon, we conducted evaluations of novel algorithms across four categories of sounds that are crucial to the experience of CI users: consonant-nucleus-consonant words (CNC) for monosyllabic word recognition, natural speech, speech in noise, and music.

#### Cochlear implant sound coding strategy

We provided a baseline software framework that incorporated a baseline sound encoding strategy, which participants could modify and enhance. Participants could choose to implement their algorithms using either Matlab or Python, or even use any programming language of their choice. The framework consisted of several component modules working together to convert audio inputs into a matrix of electrode pulses. This allowed participants to focus on developing novel algorithms for sound processing while leveraging the existing infrastructure provided by Advanced Bionics.

#### Vocoder cochlear implant simulation framework

To infer the quality of algorithms, it was necessary for participants and judges to hear the results. Since conducting actual tests on CI users was not feasible within the scope of the competition, we employed a simulated audio approach. Specifically, we used an electrodogram-based acoustic vocoder processing strategy that converted a matrix of electrode pulses into a simulated audio output resembling what a cochlear implant user would hear(11–13). This method utilized the matrix of electrode pulses for a CI as input and transformed it into an approximation of the resulting auditory experience. The vocoder strategy is an AB design with similarities to the 16-channel SPIRAL model(12). The simulated audio output served as the basis for optimizing CI processing algorithms.

#### Entry submission

Entries were validated and analyzed for errors. Once validated, entries were processed into CI simulations using a backend vocoder. Entrants could play back sounds to confirm accuracy. Entries were randomly assigned to discrete sets of audio tasks organized by category. The CI Hackathon entries were randomized alongside baseline audio outputs from the AB baseline strategy for fair comparison across four categories of audio input.

#### Judging interfaces

The research platform supports customized evaluation interfaces for assessing sound processing performance. In the CI Hackathon, we implemented different user interfaces for two rounds of judging. In the first round, judges were randomly assigned to evaluate one set of 120 audio entries, including the AB baseline strategy using a numerical scale to rate the quality and clarity of each sound. Scores were given on a scale from 1 (very poor) to 10 (excellent) (Figure 2),, and raw scores were converted into Z-scores (Figure 3) to account for any variability among judges. The mean z-score for each group of submitted entries was used for comparison and to determine the final categorical ranking. Round 2 involved pairwise comparisons (Figure 4) (between each group of submitted entries and the AB baseline strategy. Sounds from the same acoustic input were paired and presented in both orders, and judges scored each pair based on their preference. The platform was designed to accommodate large numbers of pairwise comparisons by using a matrix to split up and assign randomized pairs across judges. The total number of points earned for all grouped submissions in the same category determined the final ranking. A finite number of sets ensured that all entries were evaluated at least once, and the mean score was calculated for entries with multiple scores. The application reassigned sets for new scores only when all entries in a single category had been evaluated.

**Figure 2.**
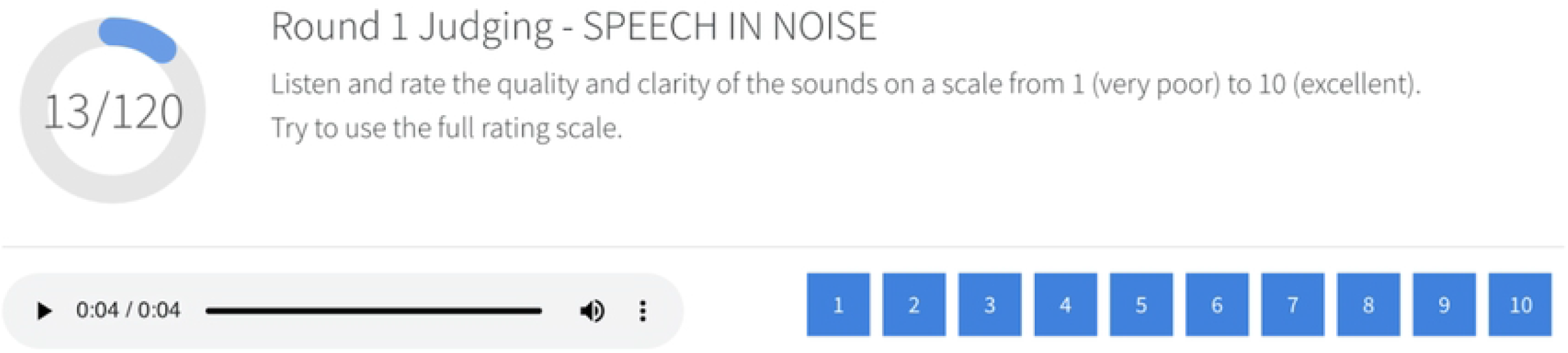
Screenshot of round 1 judging interface. The round 1 judging interface featured a progress indicator and enabled participants to subjectively rate sounds on a numerical scale. Sets of 120 sound samples within the same category were randomized and assigned to individuals and teams for judging.

**Figure 3.**
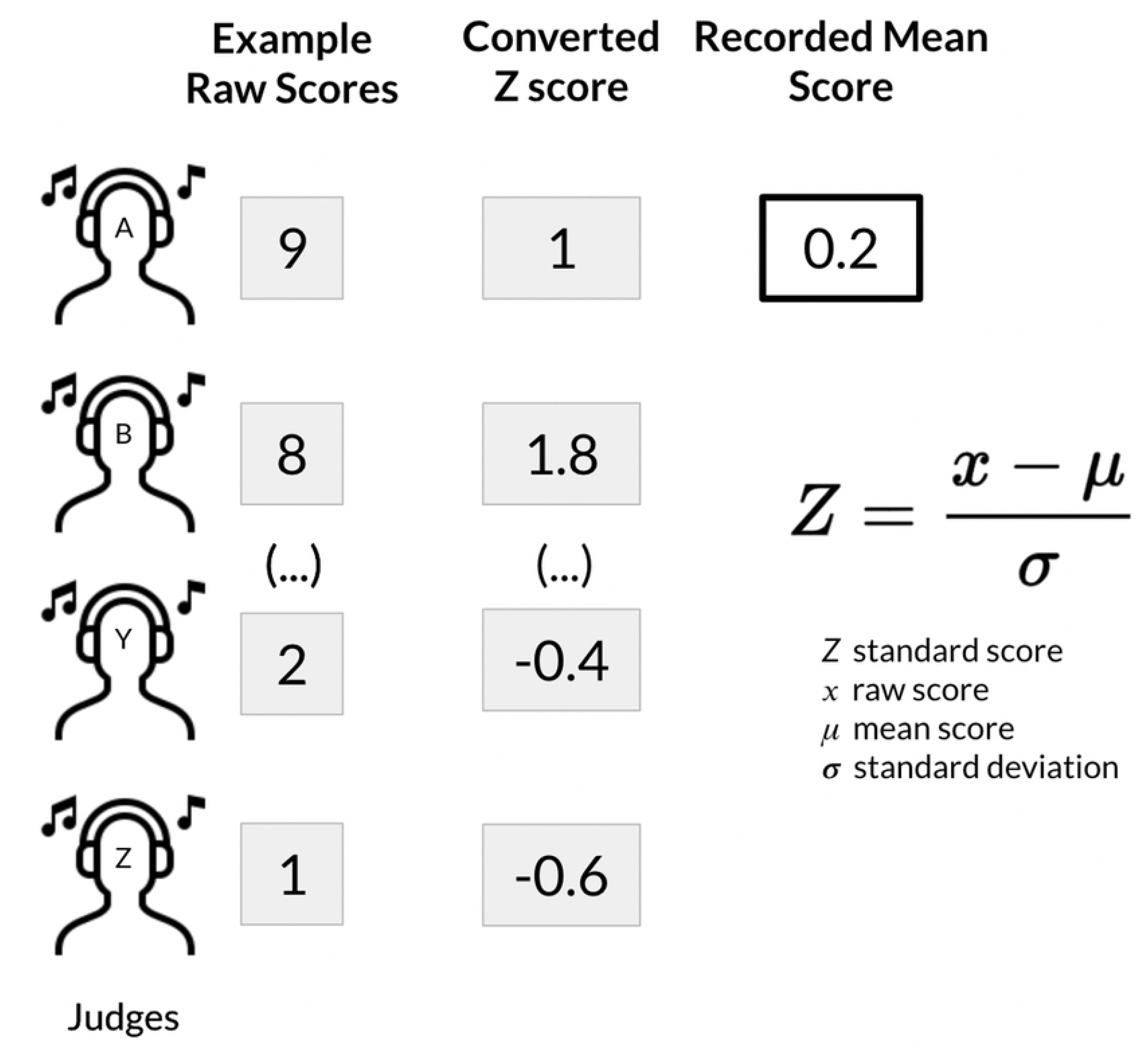
Overview of scoring for round 1. Raw scores were converted into Z-scores for each individual participant to address variability across all judges. Mean z-scores were used to rank teams in the first round of categorical ranking.

**Figure 4.**
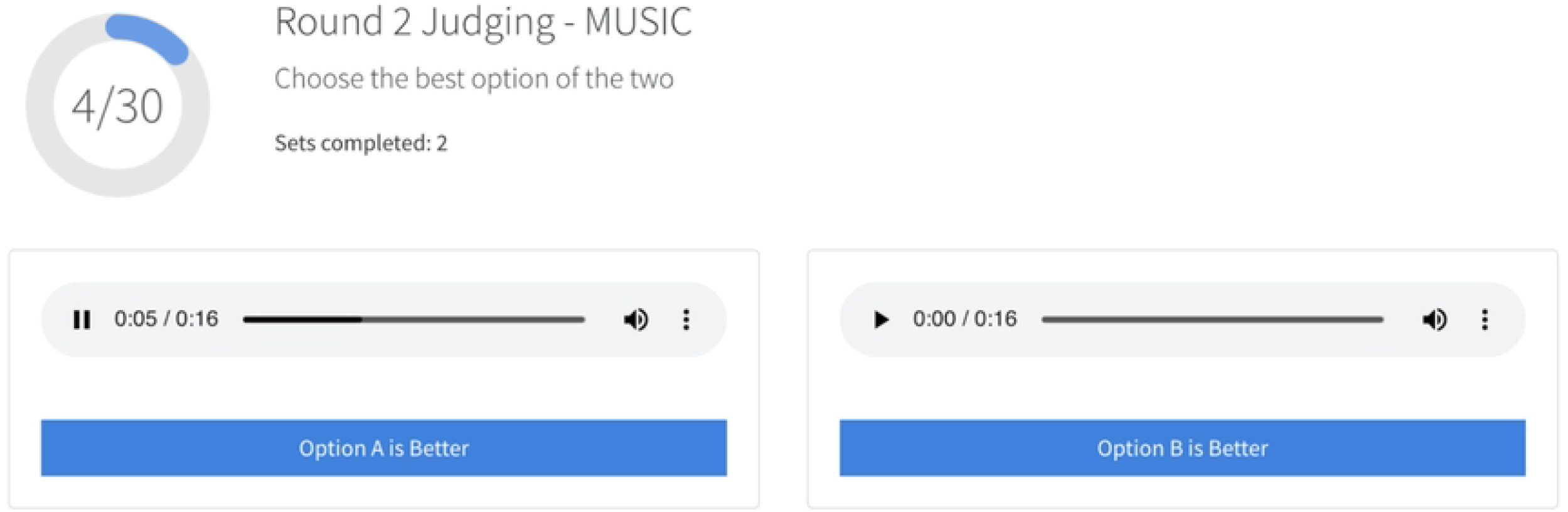
Screenshot of round 2 judging interface. The round 2 judging interface featured pairwise comparison of different sound processing strategies for audio samples from a single category.

**Figure 5.**
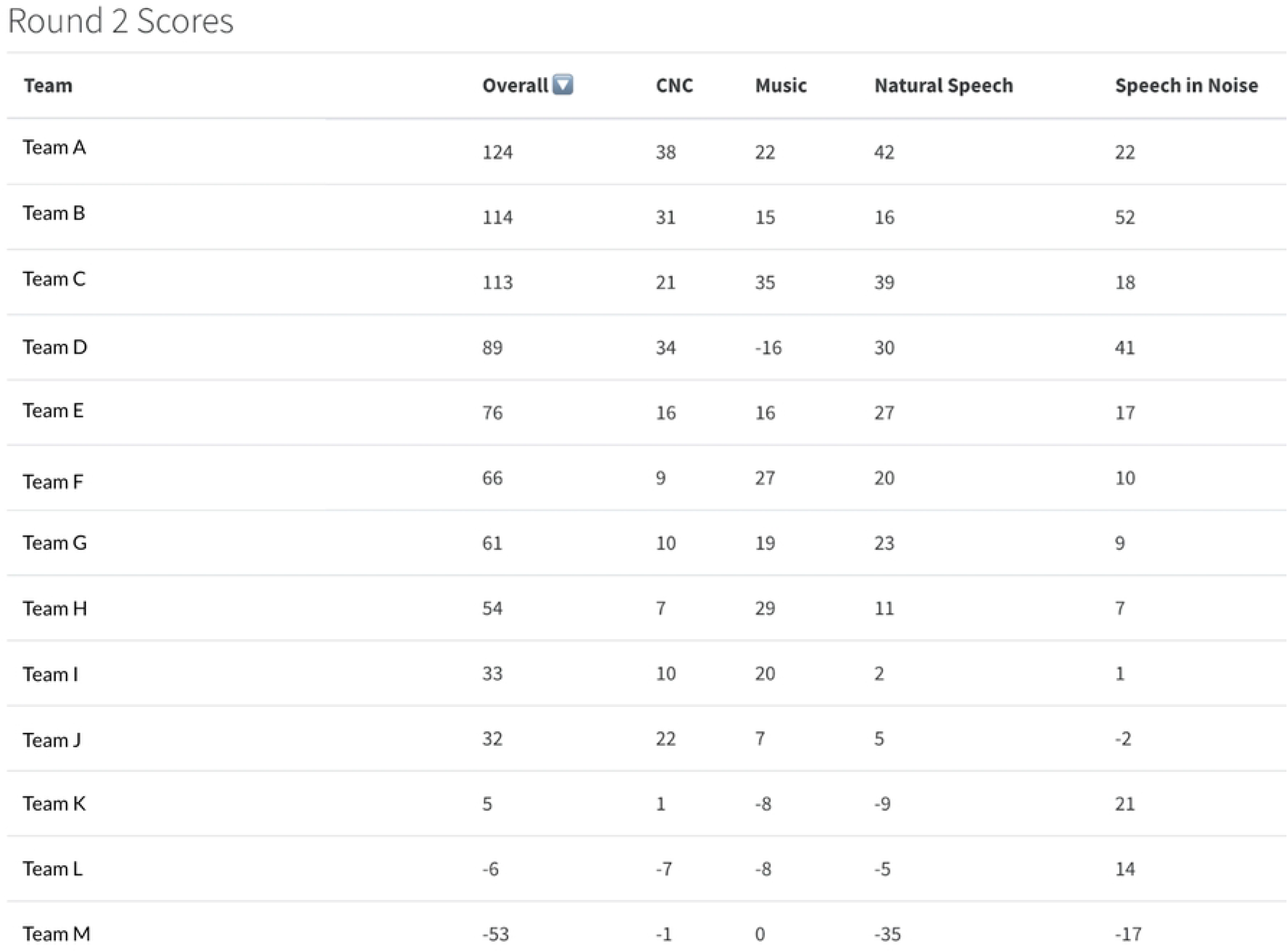
Sample output of round 2 leaderboard.

#### Scalability and Performance

To ensure the research platform’s ability to handle a substantial volume of concurrent users, we conducted an evaluation using a web-based performance testing tool. The tool generated synthetic requests, simulating user traffic, which were progressively increased in number until a threshold was reached or a failure occurred. Although AWS services are designed for scalability, performance testing can be used to identify any bottlenecks in service quotas and downstream applications. We used Artillery, an open-source tool, to generate simultaneous requests for both judging API endpoints. The median response times for 100, 1000, and 10,000 requests per hour were all less than 1 second (data not shown).

## Discussion

We introduce a web-based hearing research platform specifically designed to identify and compare different signal processing algorithms aimed at improving sound processing in cochlear implants. The platform’s design focuses on facilitating direct comparisons through acoustic simulations, providing flexibility in audio stimulus presentation and evaluation, and ensuring scalability to handle a large number of participants and entries. The virtual design enables international participation, large sample size, and enhanced internal validity. It enables researchers to conduct auditory experiments, ranging from simple to complex listening tasks, with ease of use and flexibility.

Researchers rely on a variety of platforms to evaluate the performance of experimental sound processing algorithms. Modern systems use PCs, mobile devices, and specialized hardware interfaces with onboard commercial processors. The flexibility provided by higher-level programming languages and open-source software, along with the computational power of laptops and mobile devices empowers researchers to customize various elements of the sound processing pipeline and electrode stimulation strategies. Our research platform is not limited by hardware and can conceivably be used to compare sound coding strategies more broadly and in a platform agnostic fashion.

To allow non-CI users to participate, we implemented a vocoder, acknowledging its limitations. Existing vocoders are based on signal processing and do not consider biophysical elements of sensory perception.(11) For example, CI users have limited access to pitch due to poor spectral resolution.(14) When using vocoded signals for CI simulations, numerous factors need to be considered as they impact the clarity of acoustic information. It remains uncertain whether a vocoder can truly replicate the experience of a CI user. Further work testing different algorithm variants on actual CI users will be crucial in confirming the validity of this approach.

To facilitate scalability and expedite scoring by human evaluators, we opted for short audio clips that could be easily scored using discrete numerical scales and A-B comparisons. This approach enabled standardized scoring across different types of auditory stimuli and addressed the disadvantages and challenges associated with both open and closed sets of responses.

An essential component of speech audiometry is the word recognition test (WRS), which utilizes a list of phonetically or phonemically balanced monosyllabic words known as consonant-nucleus-consonant (CNC) words. During these tests, patients are presented with words and asked to repeat them based on what they hear. The accuracy of word repetition is determined by correctly repeating all the phonemes, and the WRS score is calculated as the percentage of correctly repeated words (ranging from 0 to 100%). The use of an open set of responses in this test reflects the complexity of real-world auditory perception, allowing for a wide range of responses and identification of specific patterns that can aid in diagnosing auditory disorders. However, comparing and analyzing results across participants becomes challenging due to the unrestricted nature of open sets. To facilitate ease of scoring and comparison between novel sound processing algorithms, we opted for a numerical subjective scoring system ranging from 1 to 10. Alternatively, closed-set responses with a predetermined list of possible options can be employed. This approach enhances the efficiency, standardization, and reliability of scoring and enables quantitative analysis and comparisons between participants. However, the use of closed sets diminishes the generalizability of test results since they do not mimic real-world auditory stimuli. Ceiling and floor effects may occur, limiting the ability to obtain qualitative information. Additionally, participants may guess when uncertain, potentially impacting the accuracy of the results. Our research platform can support any modality of evaluation and scoring and is only limited by programmatic implementation.

Subjective studies involving CI recipients and experimental sound processing strategies often yield highly variable outcomes due to a combination of engineering constraints and individual patient factors. For example, challenges in music perception are commonly experienced by CI users due to technological limitations, biological factors, and difficulties in capturing the full spectral, fine-temporal, and dynamic range representation required for music comprehension.(15) Our study focused on gathering broader qualitative information rather than relying on individual perceptual tasks that are easier to quantify, but may not fully recapitulate the challenges inherent in music enjoyment.

Although the initial implementation of this platform was on vocoded speech that was scored by judges without CIs, our research platform can be directly utilized with CI users with modern processors. By temporarily deactivating sound processing on their CI processors using proprietary software, we can then deliver pre-processed audio CI simulations generated from uploaded matrixes. The CI simulations can then be streamed directly to participants’ processors via Bluetooth on a PC, tablet, or smartphone with scoring of the different simulations being done on a PC, tablet, or smartphone with the CompHEAR platform.

## Conclusion

This work demonstrates a web-based evaluation platform and accompanying tools designed for evaluating auditory CI simulations of novel sound processing algorithms. Our goal was to design and implement a standardized and scalable platform to support future hearing device research. We introduce a customizable web-based platform that utilizes performant backend technologies and user-friendly interfaces. This comprehensive platform caters to a diverse use base, including members of the public, audiologists, and clinician-scientists. Through a highly successful virtual hackathon conducted between December 2020 and January 2021, we have demonstrated the practical utility of this tool in fostering interdisciplinary collaborations and improving sound processing algorithms for CIs.

## Acknowledgements

All authors contributed to the study conception and design. The web-based judging platform was designed and implemented by KM. Material preparation, data collection, data analysis was performed by KM and LM. The first draft of the manuscript was written by KM and LM. KM had drawn the figures. Funding acquisition was performed by AJO and ADT. Supervision of the work was performed by ADT. All authors were involved and editing and review of the manuscript. All authors read and approved the final manuscript.

This project was funded by grants from Advanced Bionics and the National Institute of Deafness and Communications Disorders (R01 DC012262 awarded to AJO, R01 DC018076 to ADT). DS, PH, DA, TS, and TS are or were employees of Advanced Bionics, Inc. The authors have no other relevant competing interests to disclose.

